# Astrocytic tau inclusions lead to microglial abnormalities, but leave neuronal networks intact

**DOI:** 10.1101/2024.01.28.577642

**Authors:** Thomas Vogels, Gréta Vargová, Tomáš Hromádka

**Affiliations:** Axon Neuroscience R&D Services SE, Bratislava, Slovakia; University of California, Santa Cruz, Santa Cruz, United States; Institute of Neuroimmunology, Slovak Academy of Sciences, Bratislava, Slovakia

**Keywords:** Astrocytes, tau pathology, adeno-associated virus, tauopathies, progressive supranuclear palsy, ARTAG

## Abstract

Astrocytic tau inclusions are commonly found in the aging brain and primary tauopathies, such as progressive supranuclear palsy (PSP) and corticobasal degeneration (CBD). The functional consequences of these histopathological lesions, however, are unknown due to the lack of specific animal models. We have developed a mouse model of astrocytic tau pathology to study downstream effects on microglia and neuronal networks. We have designed an adeno-associated viral (AAV) vector expressing aggregation-prone human truncated tau (amino acids 151-391/4R) and red fluorophore mCherry in equal ratio under the astrocytic GFAP promotor. Injection of AAV-GFAP-htTau into the hippocampus of wild-type mice led to expression of human truncated tau in astrocytes and accumulation of soluble phosphorylated tau (p214, p231) but no detectable cognitive impairment. In vivo multiphoton imaging revealed alterations in microglial morphology in the vicinity of truncated tau positive astrocytes in the cortex. No alterations in firing patterns of excitatory cortical neurons surrounded by astrocytes overexpressing truncated tau were detected. These results suggest that early stages of astrocytic tau pathology lead to changes in microglial function, but not to functional impairment of neuronal networks.

## Introduction

Astrocytic tau inclusions are a prominent feature of primary tauopathies and can present in a variety of forms, such as the tufted astrocytes in progressive supranuclear palsy (PSP), astrocytic plaques in corticobasal degeneration (CBD), ramified astrocytes in Pick’s disease (PiD), globular astroglial inclusions in globular glial tauopathies (GGT), bush-like astrocytes in argyrophilic grain disease (AGD), thorn-shaped astrocytes and granular/fuzzy astrocytes in aging-related tau astrogliopathy (ARTAG) [30, 67]. ARTAG is also observed in chronic traumatic encephalopathy (CTE), Huntington’s disease, and Alzheimer’s disease (AD), and was suggested to contribute to disease symptoms in AD patients [6, 29, 48, 52, 58, 60]. Given the abundance of astrocytic tau pathology in human tauopathies and the aging brain, the functional consequences of these lesions have been understudied compared to neuronal tau pathology. Astrocytes have highly ramified morphology, which allows them to interact with virtually all surrounding cells. Astrocytes play important roles in the immune system, the vasculature, neuronal homeostasis, synaptic function, neuronal circuits, and cognition [3]. Although many of these functions were shown to be impaired in the presence of neuronal tau pathology, it remains largely unknown how astrocytic tau inclusions influence physiological functions of other cells [28, 64].

Most early transgenic mouse models with some degree of astrocytic tau pathology also featured oligodendrocytic and predominant neuronal tau pathology – thereby limiting the possibility to study astrocytic tau pathology in isolation [13, 23, 34]. Injection of brain extracts from primary tauopathy cases led to an intriguing recapitulation of disease-specific conformations and hyperphosphorylation of tau in astrocytes, but the majority of tau pathology in these models could be observed in neurons and to a lesser extent oligodendrocytes [7, 9]. Transgenic mouse models in which human tau expression was controlled under the astrocytic GFAP promotor represented significant progress in studying exclusively astrocytic tau pathology. In these mice accumulation of phosphorylated tau was detected in 30% of 12-month-old mice and 95% of 24-month-old mice, and astrocytic tau pathology was most prominent in the spinal cord [11, 17]. Efforts have also been made to model the consequences of astrocytic tau pathology using induced pluripotent stem cells, but it remains difficult to study long term pathological events in cellular models, which additionally lack the complexity of an intact nervous system [19]. Practical improvements in the development of animal models of astrocytic tau pathology in the brain parenchyma are therefore urgently needed to better understand the functional consequences of these histopathological lesions in primary tauopathies.

Here we describe an AAV-based model of astrocytic pathology that expresses human truncated tau(aa151-391/4R) and fluorophore mCherry under the astrocyte-specific GFAP promotor (AAV-GFAP-htTau). The truncated tau construct was previously shown to lead to mature neurofibrillary pathology, neuroinflammation, and associated functional deficits when expressed in neurons of mice and rats [69, 80, 81]. As tauopathies present with alterations in microglial function and neuroinflammation [68], we used AAV-GFAP-htTau to study the consequences of astrocytic tau pathology on microglial morphology. Since tau pathology drives clinical symptoms in primary tauopathies with abundant astrocytic tau pathology (e.g. PSP, CBD) [61], we also studied if expression of truncated tau in astrocytes is associated with alterations in neuronal activity in the cortex of awake mice.

## Methods

### Human brain tissue samples

Human brain samples were obtained from The Netherlands Brain Bank, Netherlands Institute for Neuroscience, Amsterdam (open access: www.brainbank.nl). All material has been collected from donors for or from whom a written informed consent for a brain autopsy and the use of the material and clinical information for research purposes had been obtained by the Netherlands Brain Bank. All human tissue samples were handled according to the standard outlines in the material transfer agreement with the Netherlands Brain Bank.

Paraffin-embedded putamen with pallidum (insula) sample was obtained from an 88 year old female patient with PSP. The post-mortem delay was 4 hours.

### Experimental animals

All animal experiments were approved by the State Veterinary and Food Committee of the Slovak Republic, and by the Ethics Committee of the Institute of Neuroimmunology, Slovak Academy of Sciences. All animals had ad libitum access to water and food and were housed under standard laboratory conditions. Mice were kept under stable conditions on a 12:12 hour light-dark cycle. The animals were anesthetized and sacrificed according to ethical guidelines to minimize pain and suffering of experimental animals. We used C57BL/6h, FVB/NJ, CX3CR1-YFP (The Jackson Laboratory, line #021160) mouse lines as specified below for each experiment. FVB/NJ mice were non-transgenic littermates from the rTg4510 line [63] and were used to minimize the need for additional breeding.

### Viral vectors

AAVDJ8-GFAP-hTau(151-391/4R)-P2A-mCherry (Vector Biolabs; AAV-GFAP-htTau) and AAVDJ8-GFAP-mCherry control (SL101522, Signagen; AAV-GFAP-Control) were both diluted to a titer of 2.75x10^12^GC/ml for intracerebral injections. Injection volumes are specified below for each individual experiment. The hTau(151-391/4R)-P2A-mCherry construct was previously used under the neuron-specific human synapsin 1 promotor [69]. The combination of the AAVDJ8 serotype and GFAP promotor has been described as driving astrocyte-specific expression of fluorophores [20, 33]. Aliquots that were thawed for experiments were kept at 4 degrees Celsius and used for a maximum of 2 weeks. AAV aliquots were blinded for all quantitative experiments and experimenters were unblinded to the identity of the injected virus only after collecting and processing the data.

### Experimental groups

*Temporal progression of astrocytic tau pathology:* To evaluate the experimental AAV, 14-month-old female FVB/NJ mice (n=4) were injected bilaterally with AAV-GFAP-htTau in the hippocampus and brains were collected after 5 days (n=1), 10 days (n=1) and 20 days (n=2). To evaluate the control AAV, 14-month-old male FVB/NJ mice (n=2) were injected with AAV-GFAP-Control in the left hippocampus and AAV-GFAP-htTau in the right hippocampus and brains were collected after 2 weeks. To evaluate the progression of tau pathology induced by AAV-GFAP-htTau, 4-9 month-old C57bl/6 of both sexes (n=20) were injected bilaterally in the hippocampus and brains were collected in monthly intervals (1-11 months post-injection, two brains per month). As a control, 6-9 month-old female C57bl/6 mice were injected with AAV-GFAP-Control in the left hippocampus and PBS in the right hippocampus (n=10), or with PBS in both hemispheres (n=10), and their brains were collected in monthly intervals (1-11 months post-injection).

#### Astrocyte specificity of viral vectors

To quantify the cell-type specificity, 2-month-old C57bl/6 wild-type mice were injected bilaterally in the hippocampus with AAV-GFAP-htTau (n=3) or AAV-GFAP-Control (n=3), and their brains were collected 2 months after injection.

#### Biochemical analysis of tau pathology

9-10 month-old female FVB/NJ mice were injected bilaterally with AAV-GFAP-htTau (n=12) or AAV-GFAP-Control (n=4) in the hippocampus. Animals were sacrificed and their brains were collected 5 months after injection.

#### Behavioral analysis

4.5-month-old male C57bl/6 wild-type mice were injected bilaterally in the hippocampus with AAV-GFAP-htTau (n=7) or AAV-GFAP-Control (n=7), and underwent behavioral testing 5 months after injection.

#### In vivo imaging of microglia

10-13 month-old CX3CR1-YFP mice were injected unilaterally in cortex with AAV-GFAP-htTau (n=6) or AAV-GFAP-Control (n=4). At 7.5-8 months after AAV injection mice were implanted with a cranial window and the morphology of cortical microglia were imaged in-vivo.

#### In vivo imaging of neuronal activity

1-2 month-old C57bl/6 wild-type mice were injected with AAV-GFAP-htTau (n=3) or AAV-GFAP-Control (n=3). At 4.5-6 months after AAV injection mice were injected with an AAV expressing calcium indicator GCaMP6f, implanted with a cranial window, and the activity of cortical neurons was imaged in-vivo.

### Intracerebral AAV injections into hippocampus

Intracerebral injections into the hippocampus were performed as previously described [69]. Anesthesia was induced by 3% isoflurane, after which the mice were mounted in a stereotaxic apparatus (Robot Stereotaxic, Neurostar). Isoflurane (1%) was used to maintain optimal levels of anesthesia. The local anesthetic lidocaine was injected subcutaneously under the scalp to minimize pain at the site of the incision. For each AAV a dedicated 10μl Hammilton syringe (SGE) was used, and the needle was cleansed internally by repeatedly aspirating 70% ethanol followed by sterile saline between injections. The following injection coordinates were used: AP: -2.5 mm and L: ±2 mm. The animals were first injected with 1 μl at a depth of -2.2 mm from brain surface (100 nl/min + 5 min wait-time). The needle was then carefully withdrawn by 600 μm to 1.6mm depth and the same injection protocol was repeated. After the surgery, the wound was closed with tissue adhesive (Vetbond), and animals were allowed to recover on a heating pad. Rimadyl (5mg/kg, intraperitoneal) was applied for post-surgical analgesia.

### Cortical AAV injections

Cortical AAV injections were performed in transgenic CX3CR1-YFP or wild-type mice, to image microglia and neuronal activity (GCaMP6f) respectively. The surgical steps to prepare the mouse for injection were identical to the described protocol for injections into the hippocampus. However, instead of drilling through the skull, the skull was thinned as much as possible. The tip of a sterile insulin needle was used to peel away the remaining thin layer of skull to expose the surface of the brain. Glass micropipettes (Drummond Scientific Company, ∼15 μm tip diameter) were used to inject the AAV into the cortex with minimal damage to neural tissue and minimal AAV leakage to the surface.

AAV-GFAP-htTau or AAV-GFAP-Control were loaded into the micropipette by pipetting a drop of AAV onto a small piece of parafilm. Under visual control using a surgical microscope, the required volume of AAV was slowly aspirated into the pipette. The tip of the glass pipette was then lowered through the craniotomy until it penetrated the dura. A 5ml syringe was attached to the end of the injection tubing and positive pressure was slowly applied to inject the virus. The speed and volume of the injection (20–30 nl/min, 500 nl total) were controlled by visually monitoring the movement of the virus solution in the pipette shaft across the calibration marks. The pipette was first lowered to 500-600 μm below the surface of the brain to inject 250 nl of AAV, and then raised by 300 μm to inject another 250 nl of AAV. After waiting for 2-3 min, the pipette was withdrawn slowly to avoid backflow of the virus solution to the surface. The skin was carefully closed as described above.

Calcium indicator GCaMP6f (AAV1-Synapsin-GCaMP6f-WPRE-SV40, Addgene 100837-AAV1; 6.52 ×10^12^GC/ml) was injected using the same procedure at 4.5-5 months after injection of AAV-GFAP-htTau or AAV-GFAP-Control. Injections (2-3 injections of 100 μl each per mouse, 200-250 μm deep, 50 nl/minute) were carefully centered around the previous AAV injection site. A cranial window was implanted immediately after the injection of the calcium indicator as described below.

### Cranial window implantation

Before the surgery, the mouse was injected with dexamethasone (2 mg/kg) to reduce potential swelling and inflammation of the brain. The mouse was then prepared as described previously for AAV injections. However, instead of making an incision, a piece of skin above the target area was removed, underlying bone was cleaned and carefully dried, and a custom-made headbar was attached to the bone using dental acrylic (Duracryl Plus). A circular craniotomy was made using a round 3mm diameter biopsy puncher(Kruuse). A 3mm glass coverslip was placed on top of the dura inside the craniotomy and gently pressed down with a custom-made holder. Cyanoacrylate glue (Krazy Glue) was then carefully applied to the edges of the craniotomy to permanently fix the window. All exposed skull and the edge of the window were covered with dental acrylic for additional stability. The mouse was the injected with analgesic (carprofen, Rimadyl 5mg/kg) and antibiotic (enrofloxacin, Baytril, 5mg/kg) and allowed to recover on a heating pad. After recovery, the mouse was injected daily with a mixture of carprofen and enrofloxacin for 7 days, and dexamethasone for the first 3 days, as described previously [50]. Calcium imaging was performed 2-3 weeks after cranial window implantation. Microglia were imaged 3-4 weeks after cranial window implantation as described previously [77].

### Tissue collection and preparation

The collection of brain samples was performed as described previously [69]. An intraperitoneal administration of zolazepam/tiletamine (Zoletil 100, 100 mg/kg, Virbac) and xylazine (Xylariem, 20 mg/ml, Ecupharn N.V.) in a ratio of 1:1.7 was used to deeply anesthetize the mouse. For experiments that only required tissue for histological analysis, mice were transcardially perfused with PBS containing 2% heparin until the blood was removed, followed by 3 minutes perfusion with cold PBS containing 4% paraformaldehyde for fixation, after which brain samples were carefully removed for histological analysis. For experiments where brain tissue was dissected out for biochemical analysis, animals were only perfused with PBS with 2% heparin for 3 minutes. The hippocampus was dissected from one hemisphere, frozen in liquid nitrogen, and stored at -80 degrees Celsius for biochemical analysis. The opposite hemisphere was used for histological analysis. Histological samples were post-fixed overnight in PBS containing 4% paraformaldehyde at 4 degrees Celsius. The following day, brains were rinsed with PBS and cryoprotected using PBS containing 30% sucrose at 4 degrees Celsius until the brain samples sank completely. Brain samples were then mounted in cryomedium blocks (Leica, Wetzlar, Germany) and frozen at -20 degrees Celsius for later sectioning with a cryomicrotome (Leica CM 1850, Leica, Wetzlar, Germany).

### Immunohistochemistry

Immunohistochemistry was performed at room temperature as described previously [69]. Two hours of immersion in 4% PFA was used for post-fixation of the cryosections (40 µm). After washing in PBS, antigen retrieval was performed by immersing the sections in ice-cold 80% formic acid for 40 seconds. Immersion in 1% hydrogen peroxide was used to block endogenous peroxidase activity. After washing, sections were permeabilized by immersion in PBS containing 0.3% triton for 10 minutes. Blocking was performed by immersing sections in APTUM section block (Diagnostic Technology) for 1 hour, after which they were incubated overnight in primary antibodies dissolved in PBS at 4° Celsius (Table 1). Final staining was performed using the Vectastain ABC Kit (Vector Laboratories, CA, USA). After thorough washing, sections were incubated in secondary antibodies for two hours and subsequently incubated in ABC solution for 1 hour. The Vector VIP kit (Vector Laboratories) was used to develop the final signal. PBS-washed sections were then mounted on glass microscope slides and left to dry for at least 2 hours. Slides were then immersed in PBS (10 min), 70% ethanol (5 min), 96.5% ethanol (5 min), isopropylalcohol (5 min), xylene (2 x 5 min) and coverslipped using Entallan mounting medium (Merck). Samples were imaged using an Olympus BX51 microscope equipped with an Olympus DP27 digital camera (Olympus Microscope Solutions).

**Table 1.** Primary antibodies used in this study.

| Antibody/epitope | Host species | Dilution | Source; antibody ID |
| --- | --- | --- | --- |
| HT7 (hTau aa159-163) | Mouse | 1:500 | Thermo Scientific (IL, USA); MN1000 |
| DC18 (hTau aa168–181) | Mouse | 1:100* | Axon Neuroscience (Bratislava, Slovakia) |
| AT270 (tau pT181) | Mouse | 1:500 | Thermo Scientific (IL, USA); MN1050 |
| AT8 (tau pS202/pT205) | Mouse | 1:500 | Thermo Scientific (IL, USA); MN1020 |
| Tau pT212 | Rabbit | 1:500 | Thermo Scientific (IL, USA); 44-740G |
| Tau pS214 | Rabbit | 1:500 | Thermo Scientific (IL, USA); 44-742G |
| DC217 (tau pT217) | Mouse | 1:100 | Axon Neuroscience (Bratislava, Slovakia) |
| AT180 (tau pT231) | Mouse | 1:500 | Thermo Scientific (IL, USA); MN1040 |
| DC25 (pan-tau aa347–353) | Mouse | 1:1* | Axon Neuroscience (Bratislava, Slovakia) |
| Neun (neuronal marker) | Rabbit | 1:1000 | Abcam; ab104225 |
| CNP (oligodendrocyte marker) | Mouse | 1:500 | Merck; MAB1580 |
| GFAP (astrocyte marker) | Rabbit | 1:1000 | Abcam; ab7260 |
| IBA1 (microglia marker) | Rabbit | 1:500 | Wako (Tokyo, Japan); 019-19741 |

*\*Hybridoma supernatant*

The protocol for paraffin embedded sections was identical, with the exception that slides were first deparaffinized by immersion in xylene (2 x 5 min), isopropylalcohol (5 min), 96.5% ethanol (5 min), 70% ethanol (5 min), and Millipore water (5 min). For comparison of sections from AAV-GFAP-htTau injected animals and PSP and CBD patients, all sectioning (8 µm thickness) and staining was performed in the same session and under the same experimental conditions.

### Immunofluorescence on free floating sections

Immunofluorescence staining was performed at room temperature (unless specified otherwise) as described previously [69]. Permeabilization of cryosections (40 µm) was performed by immersing them in PBS containing 0.3% triton. APTUM section block (Diagnostic Technology) was used for one hour to block the sections, after which they were incubated overnight in primary antibodies dissolved in PBS at 4° Celsius (Table 1). The following day, sections were incubated in secondary antibodies in PBS. Vectashield mounting medium containing DAPI (Vector Laboratories) was used to coverslip the slides. A Zeiss 710 Laser Scanning Confocal microscope (Zeiss, Jena, Germany) using 10×, 20×, 40× and 63× objectives was used to image the sections. The gain of photo-multiplier tubes was optimized for each imaging session, while keeping the laser power constant, and was kept the same for all slides when AAV-GFAP-Tau and AAV-GFAP-control were compared.

### Sarkosyl extraction

Sarkosyl isolation was performed as described previously [69]. Hippocampi from mice injected with AAV-GFAP-htTau or AAV-GFAP-Control (5 months post-injection in the hippocampus) were pooled (4 hippocampi/tube) and homogenized in 10 volumes of ice-cold homogenization buffer (20 mM Tris, 0.8M NaCl, 1 mM EGTA, 1 mM EDTA, and 10% sucrose) supplemented with protease (Complete, EDTA free, Roche Diagnostics, USA) and phosphatase inhibitors (1 mM Sodium orthovanadate, 20 mM NaF), followed by centrifugation at 20,000 g for 20 min. Supernatant was adjusted to 1% (w/v) N-lauroylsarcosine and incubated for 1 hour at room temperature with stirring. After incubation supernatant was centrifuged at 100,000 g for 1.5 h at 25°C. Pellets were re-suspended gently in 1/50 of initial homogenization volume in 1xSDS sample loading buffer.

### Western blot

Sarkosyl insoluble proteins were separated using 12% SDS-PAGE gels and transferred to nitrocellulose membrane as described previously [81]. The membrane was incubated with primary antibodies diluted in 5% non-fat dry milk for 1 h or overnight at 4°C or according to manufacturer’s instructions. Blots were developed using enhanced chemiluminescence detection system (SuperSignal West Pico chemiluminiscence substrate, Thermo Scientific, USA).

### Behavioral experiments

Movement of animals in all behavioral experiments was recorded using a camera (PointGrey Flea3, FL3-U3-13S2M-CS) mounted on top of behavioral enclosures. Bonsai software was used to track the position of the mouse in the enclosure/maze [38].

*Open field and novel object recognition.* Only male mice were used for behavioral experiments to reduce variation in mouse behavior caused by the menstrual cycle. Mice were habituated to the behavior room for 7 days before the start of the experiment and were handled daily to reduce stress during the experiments. On experiment day 1, mice were placed in a custom-made behavioral enclosure (45 cm × 45 cm × 45 cm box) opened on top and were allowed to move freely for 10 minutes. After each experimental session, the mouse was placed back to its home cage. To minimize olfactory cues, the box was cleaned with 70% ethanol, which was left to evaporate before placing the next animal in the cage.

On experiment day 3, two identical clean objects were placed in pre-selected locations in the behavioral enclosure and the mouse could explore them freely for 10 minutes (*familiarization phase*). Both the objects and the box were then cleaned as described above. After 4 hours, one novel object and one object identical to the previous session were positioned in the same box, and the mouse was tested by exploring the objects for 10 min (*test phase*). Object locations in both the familiarization and testing phases were randomized (left/right) for each mouse and group.

Total distance traveled, speed, and open field coverage were evaluated for each mouse and each open field session. Novel object recognition was assessed using *novelty preference* computed as (time spent exploring novel object)/(total time exploring objects)*100.

*Y-maze spontaneous alternations test.* The Y-shaped maze consisted of three identical arms (36 × 8 × 20 cm, length × width × height) spaced 120° apart, opened on top. The mouse was placed in end of one arm and allowed to explore the Y-maze freely for 10 minutes. Spontaneous alternations, i.e. the mouse entering a different arm of the maze in each of three consecutive arm entries, were evaluated by computing *spontaneous alternation %* = (number of spontaneous alternations)/(total number of arm entries-2)*100.

### In vivo 2-photon imaging of microglia

The mouse was anesthetized with injectable anesthetics as described above. No isoflurane was used as it could affect microglial surveillance and ramification *in vivo* [40]. The cranial window was cleaned with 70% ethanol and the mouse was placed on a heating pad to maintain a physiological body temperature. Clear immersion gel was applied between the cranial window and the objective (16×, 0.8 NA, Nikon).

Two-photon imaging was performed using a multiphoton microscope (Sutter Moveable Objective Microscope; MOM) equipped with 8kHz resonant scanner and a tunable femtosecond laser (Insight DeepSee+, Spectra Physics), and controlled by ScanImage 5 (Vidrio Technologies) [56]. In each mouse, images (512×512 pixels, approx. 150×150 μm)were acquired at 30Hz frame rate in four non-overlapping regions during a single two-hour long imaging session. Z-stacks of 40-55 μm (0.6μm steps, 30 frames/slice) were obtainedin two-minute intervals, for a total of 20 minutes per region (10 z-stacks/region). The laser power was set to 30-55 mW (measured during scanning). Microglia z-stacks were imaged at 930 nm and one corresponding z-stack in each field of view was obtained at 1040 nm to image the mCherry-positive astrocytes surrounding the microglia.

Slices in stacks were averaged and filters were applied using the Despeckle and Median Filter plugins in Fiji as described [32]. The soma size was estimated by measuring the area of the soma maximum projection in a z-stack. Only those somas that did not extend beyond the imaged region, and did not overlap with other somas or large bright processes which could confound interpretation, were measured. Care was taken not to include the beginning of large processes into soma measurements. At least ten somas per mouse were analyzed. Process length was measured semi-automatically using the Simple Neurite Tracer plugin in Fiji [37]. Only clearly visible processes that did not extend over the edges of any of the 3 dimensions were included. Further, we did include processes that overlapped such that the cell of origin could not be determined. Processes from at least 5 cells per mouse were included.

### In vivo 2-photon imaging of neuronal activity

Two-photon calcium imaging was performed in awake head-fixed mice sitting in a fixed tube platform to minimize movement. To reduce stress caused by head-fixation, mice were habituated to the experimental apparatus before performing the experiment. For imaging GCaMP6s labeled neurons we used 930nm wavelength, whereas mCherry was acquired at 1040nm. Cortical layer 2/3 spontaneous Ca2+ fluorescence signals were recorded daily, across multiple fields of views, and each was recorded for at least 5 minutes.

Raw calcium data were extracted using Suite2P [54], which performs several processing stages, including registration of the frames to account for brain motion artifacts. Regions of interests (ROIs) were selected manually using Suite2P graphical user interface (GUI), which separates somata from dentritic processes based on morphology. For analysis we used custom Matlab scripts (Mathworks). After registration of movies, background subtraction, and ROI selection, calcium signals for each ROI were calculated as changes in fluorescence defined as ΔF/F = (F − F_0_)/F_0_, where the baseline F_0_ was calculated by taking a short sliding window over 1000 frames (33s) and taken 20^th^ percentile of values of that window. Frequency of Ca2+ transients were analyzed by first generating Z-scored ROI fluorescence traces for each FOV. Ca2+ transients were identified as changes in relative fluorescence that were two standard deviations above the baseline for 10 consecutive frames (330 ms). Afterwards, neurons were classified based on the frequency of Ca2+ transients during a one-minute time-course.

### Statistical analysis

For all experiments, unpaired two-tailed Student’s t-tests were used to test for differences in means of respective parameters between groups of animals infected with AAV-GFAP-htTau or AAV-GFAP-Control. No corrections for multiple comparisons were necessary.

## Results

### Inducing selective expression of human truncated tau in astrocytes

To induce selective astrocytic expression of human truncated tau, we have designed an AAV (serotype DJ/8) vector capable of inducing truncated tau-specific histopathological changes selectively in astrocytes. AAVDJ8-GFAP-hTau(151-391/4R)-P2A-mCherry (AAV-GFAP-htTau) co-expresses non-mutated human truncated tau(151-391/4R) and mCherry red fluorescent protein in a 1:1 ratio. **(Figure 1A)**. A related AAV construct has been used previously to induce neuron-specific tau pathology [69]. For *in vivo* transduction of human truncated tau and fluorophore mCherry in astrocytes, we performed stereotaxic injections of AAV-GFAP-htTau, AAV-GFAP-Control (a control virus), or PBS in the hippocampus of wild-type mice (**Figure 1A)**. Mice between 4-14 months of age (see Methods for details) were injected with 2 μl of AAV (10^12^ GC/ml) per hemisphere. Three weeks after injection of AAV-GFAP-htTau, we detected widespread accumulation of mCherry-positive signal in all subregions of the hippocampus **(Figure 1B).**

**Figure 1.**
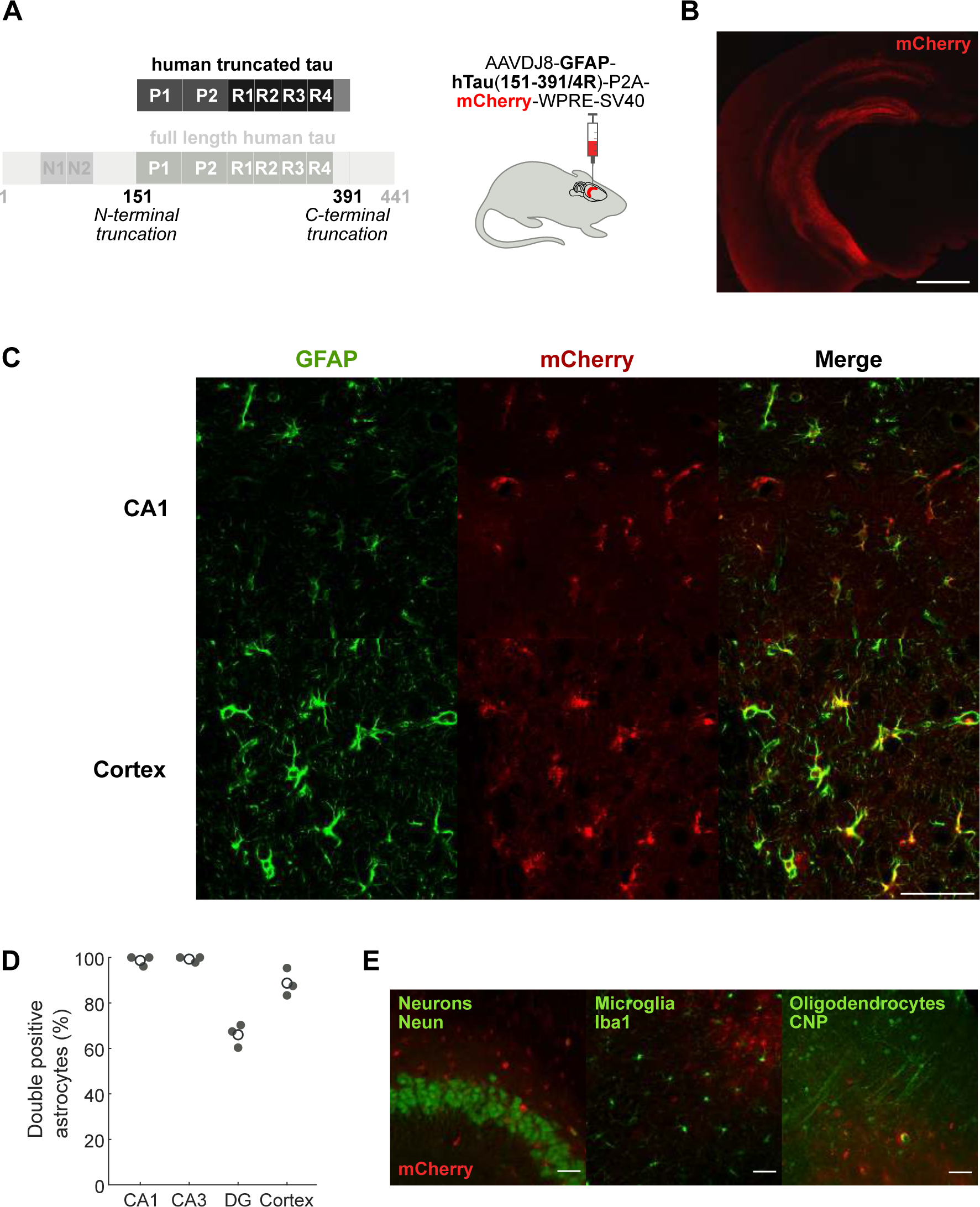
Injection of AAV-GFAP-htTau led to astrocyte-specific expression of human truncated tau. **(A)** AAV-GFAP-htTau delivering the double truncated human tau(151-391/4R) construct (left) was injected into the hippocampus. **(B)** Distribution of cells positive for mCherry (red)–expressed in a 1:1 ratio with htTau–along all subregions of the hippocampus three weeks after injection of AAV-GFAP-htTAu. Scale bar shows 1 mm. **(C)** Co-localization of mCherry (red), indicating expression of htTau, and astrocytic marker GFAP (green) in CA1 and the cortex overlaying the hippocampus. Scale bar shows 50 μm. **(D)** Quantification of the co-localization of mCherry with GFAP in CA1, CA3, dentate gyrus (DG), and the cortex overlaying the hippocampus. Percentage of double-positive cells was quantified in two slices for each mouse (n=3 mice), empty circles show means. **(E)** Expression of mCherry (red) was rarely detected in neurons (NeuN), and was not detected in microglia (IBA1) and oligodendrocytes (CNP). Areas shown are dentate gyrus (NeuN) and CA3 of the hippocampus (IBA1, CNP). Scale bars show 50 μm.

The combination of the AAVDJ8 serotype and GFAP promotor can drive astrocyte-specific expression of fluorophores (e.g. mCherry) [20, 33]. We observed strong co-localization of astrocytic marker GFAP with mCherry-expressing cells in the CA1 (98.7%, n=3 mice, 2 slices/mouse; total of 54, 76, 51 mCherry-expressing cells counted per mouse) and CA3 (99.3%, n=3 mice, 2 slices/mouse; 57, 51, 46 mCherry+ cells per mouse) regions of the hippocampus and overlying cortex (88.7%, n=3 mice, 2 slices/mouse; 40, 43, 48 mCherry+ cells per mouse) (**Figure 1C**, **Figure 1D**). Selectivity was lower in dentate gyrus (66%, n=3 mice, 2 slices/mouse; 43, 74, 58 mCherry+ cells per mouse) and weak mCherry signal was observed in neuronal-like cell bodies in neuron-dense areas such as the granular layer of the dentate gyrus (**Figure 1B**, **Figure 1D**). We observed occasional co-localization of mCherry with mature neuronal marker NeuN in the dentate gyrus (**Figure 1E**). No co-localization was observed with oligodendrocytes (CNP) or cells of myeloid origin such as microglia (IBA1) (**Figure 1E**). This indicates that in some regions there might be weak GFAP-driven expression in neurons, as has been described previously for different GFAP promotor constructs [78]. Taken together, these results suggest that AAV-GFAP-htTau drives expression of mCherry with high astrocytic selectivity.

### Accumulation of human truncated tau in astrocytes

To confirm the presence of human truncated tau in astrocytes, we used human-tau specific antibody HT7 on tissue of mice 2 months after injection of AAV-GFAP-htTau or control. Co-localization of mCherry with human tau in cells with astrocytic morphology observed in mice injected with AAV-GFAP-htTau, but not in mice injected with AAV-GFAP-Control (**Figure 2A)**. Human tau-positive astrocytes were also observed with human tau-specific antibody DC18 (not shown). The presence of a weak band of sarkosyl insoluble tau five months after injection suggests that most of the astrocytic truncated tau was soluble with only a limited amount of aggregated tau in astrocytes (**Figure 2B).**

**Figure 2.**
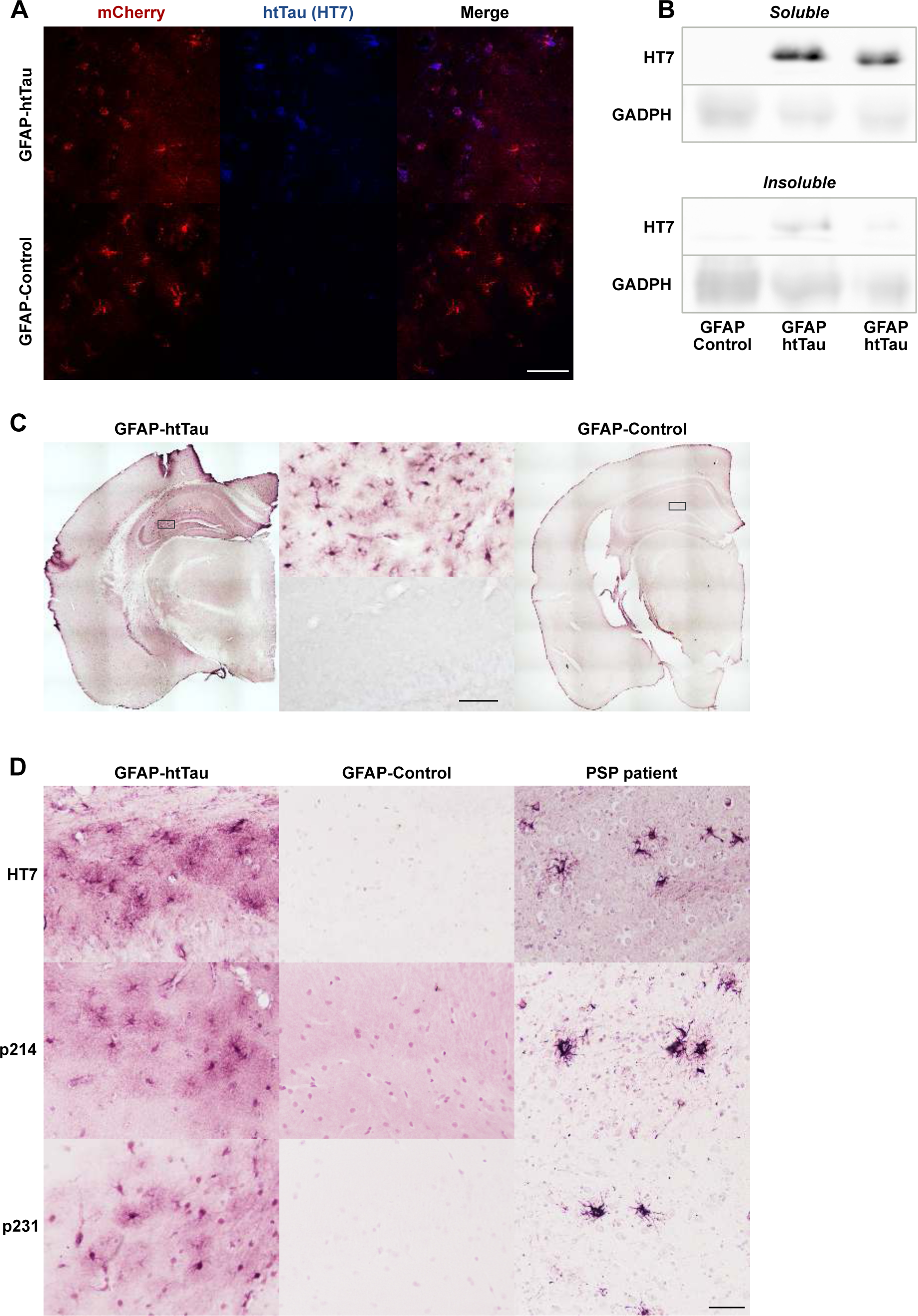
Human truncated tau accumulated in astrocytes after injection of AAV-GFAP-htTau. **(A)** Immunofluorescence staining on free floating sections showing the presence of human tau (HT7) in mCherry-positive astrocytes at 2 months after AAV injection in hippocampus. Scale bar shows 50 μm. **(B)** Western blot of both soluble and insoluble fractions of mice injected with AAV-GFAP-htTau or AAV-GFAP-Control. Detected with pan-tau antibody DC25. **(C)** Immunohistochemical staining on free floating sections showing the hippocampal distribution of human tau (HT7) at 2 months after AAV injection. Scale bar shows 50 μm. **(D)** Immunohistochemical stainings (HT7, p214, p231) on paraffin embedded sections of cortical slices from mice injected with AAF-GFAP-htTau or AAV-GFAP-Control at 8 months after injection, and from a patient with PSP (right column).

Widespread expression of human tau in cells with astrocytic morphology was observed at 2 months after injection of AAV-GFAP-htTau, but not in control animals (**Figure 2C**). The human tau positive astrocytes at 2 months after injection showed strong human tau signal in the soma and primary processes. An additional cloud of diffuse staining surrounding all human tau-positive astrocytes sug-gested that the human tau was also abundantly present in the highly ramified astrocytic endfeet (**Figure 2C**).

PSP patients accumulate 4R tau inclusions in astrocytes, known as tufted astrocytes [24]. To compare astrocytes in our model with astrocytes of patients with PSP, we stained paraffin-embedded sections from mice and a PSP patient under the same conditions in the same session. Tau mRNA and protein are not detectable in astrocytes in the healthy brain [18, 27, 49, 79], but both mice injected with AAV-GFAP-htTau, and patients with PSP accumulated human tau in astrocytes (**Figure 2D**). In astrocytes of AAV-GFAP-htTau injected animals we detected strong somatic signal from human tau with a surrounding cloud of signal likely derived from astrocytic processes (**Figure 2D**), analogous to stainings in free floating sections (**Figure 2C**). The intensity of signal detected in the soma and processes appeared comparable between sections from AAV-GFAP-htTau injected animals and human brain sections from a PSP patient (**Figure 2D**).

### Accumulation of phosphorylated tau in astrocytes

Abundant presence of hyperphosphorylated tau is characteristic for astrocytic tau pathologies in PSP and other primary tauopathies [24]. Interestingly, we observed positive staining with phospho-tau antibodies p214 and p231 at 8 months after AAV injection **(Figure 2D)**, as well as at earlier and later timepoints (5-9 months after injection). Similar to the human tau stainings, the somatic signal in the astrocytes mainly resembled the tufted astrocytes in PSP. Human tau signal in the finer processes could be observed in the PSP patient, but was more pronounced in animals injected with AAV-GFAP-htTau **(Figure 2D)**. We also observed a limited number of astrocytes positive for p181, p212, and p217, but not AT8 (pS202/pT205) (not shown). In contrast to an analogous neuronal model [69], we did not observe methoxy-X04 positive cells (not shown). These results suggest that astrocytic tau pathology in AAV-injected mice represents a relatively early stage, with pronounced expression of phosphorylated truncated tau in somas and processes of astrocytes.

### Early astrocytic tau pathology was not associated with detectable cognitive changes

To investigate whether astrocytic tau pathology leads to cognitive impairment, we assessed spatial recognition and memory of AAV-injected mice at five months after the injection. **(Figure 3)**. We did not detect differences in open field coverage and total distance traveled between mice injected with AAV-GFAP-htTau or AAV-GFAP-Control (**Figure 3A**, Field coverage: GFAP-htTau mean=2.57, CI=2.28–2.77; GFAP-Control mean=2.77, CI=2.58–2.99. Total distance: GFAP-htTau mean=33.69, CI= 29.95–41.24; GFAP-Control mean=31.09, CI=28.76–35.65). We next tested the two groups of AAV-injected mice in Y-maze spontaneous alternations test and in novel object recognition test with a testing interval of four hours. Similarly, we detected no clear differences between the two groups of mice in Y-maze spontaneous alternations **(Figure 3B**, GFAP-htTau mean=36.43, CI=32.43–41.70; GFAP-Control mean= 28.14, CI=19.17–37.43), novelty preference **(Figure 3C**, GFAP-htTau mean=66.98, CI=52.64–78.91; GFAP-Control mean=66.66, CI=19.17–37.43). These results suggest that astrocytic tau pathology in AAV-GFAP-htTau injected mice, corresponding to the early stages of human astrocytic tau pathology, did not lead to observable cognitive decline.

**Figure 3.**
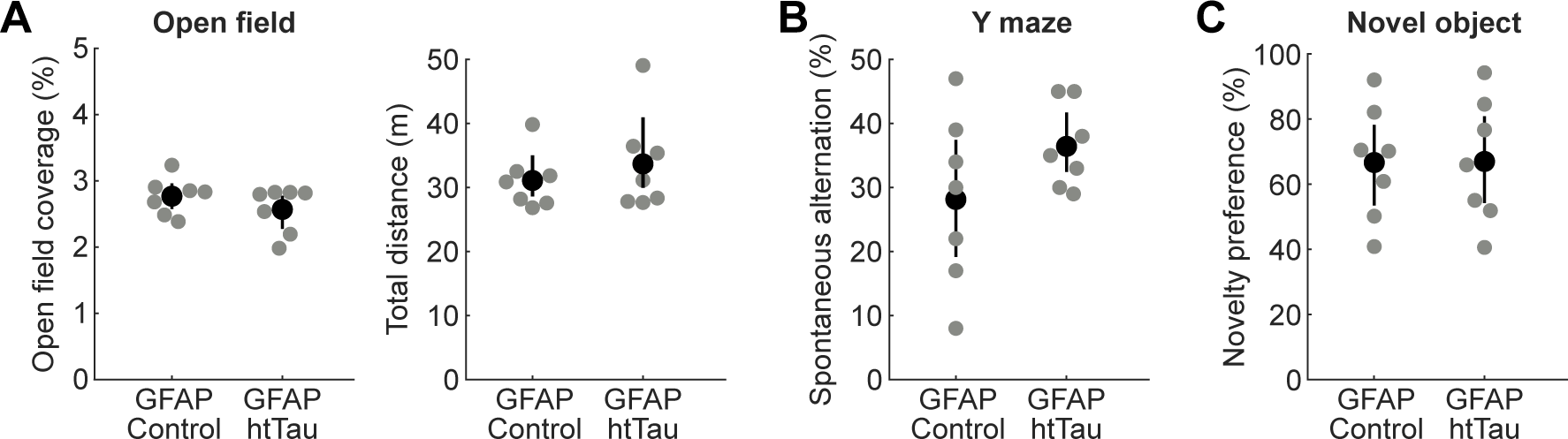
Astrocyte-specific tau pathology did not result in detectable behavioral impairments. **(A)** Mice injected with AAV-GFAP-htTau (n=7) or AAV-GFAP-Control (n=7) explored the same percentage of area and covered the same total distance in a behavioral arena during open-field testing. Field coverage: GFAP-htTau mean=2.57, CI=2.28–2.77; GFAP-Control mean=2.77, CI=2.58–2.99. Total distance: GFAP-htTau mean=33.69, CI= 29.95–41.24; GFAP-Control mean=31.09, CI=28.76–35.65. **(B)** Percentage of spontaneous alternations in Y-maze was similar in AAV-GFAP-htTau (n=7) and AAV-GFAP-Control (n=7) mice. GFAP-htTau mean=36.43, CI=32.43–41.70; GFAP-Control mean= 28.14, CI=19.17–37.43. **(C)** The overall preference for novel objects was the same in mice injected with AAV-GFAP-htTau (n=7) or AAV-GFAP-Control (n=7). GFAP-htTau mean=66.98, CI=52.64–78.91; GFAP-Control mean=66.66, CI=19.17–37.43. Gray circles in all panels correspond to individual mice, black circles and error bars show means and bootstrap estimates of 95% confidence interval of the mean.

### Astrocytic tau pathology was associated with changes in microglial morphology

To test whether astrocytic tau pathology is associated with alterations in microglial morphology and loss of homeostatic microglial behavior we injected AAV-GFAP-htTau or AAV-GFAP-Control in the cortex of Cx3cr1-YFP mice–a line with fluorescently labelled microglia–and imaged the mice after 7-8 months **(Figure 4A)**. In vivo imaging revealed that mCherry-positive astrocytes surrounded cortical microglia **(Figure 4B).** Transduction of the cortex and sustained expression of human truncated tau was confirmed in brain slices **(Figure 4C-D)**.

**Figure 4.**
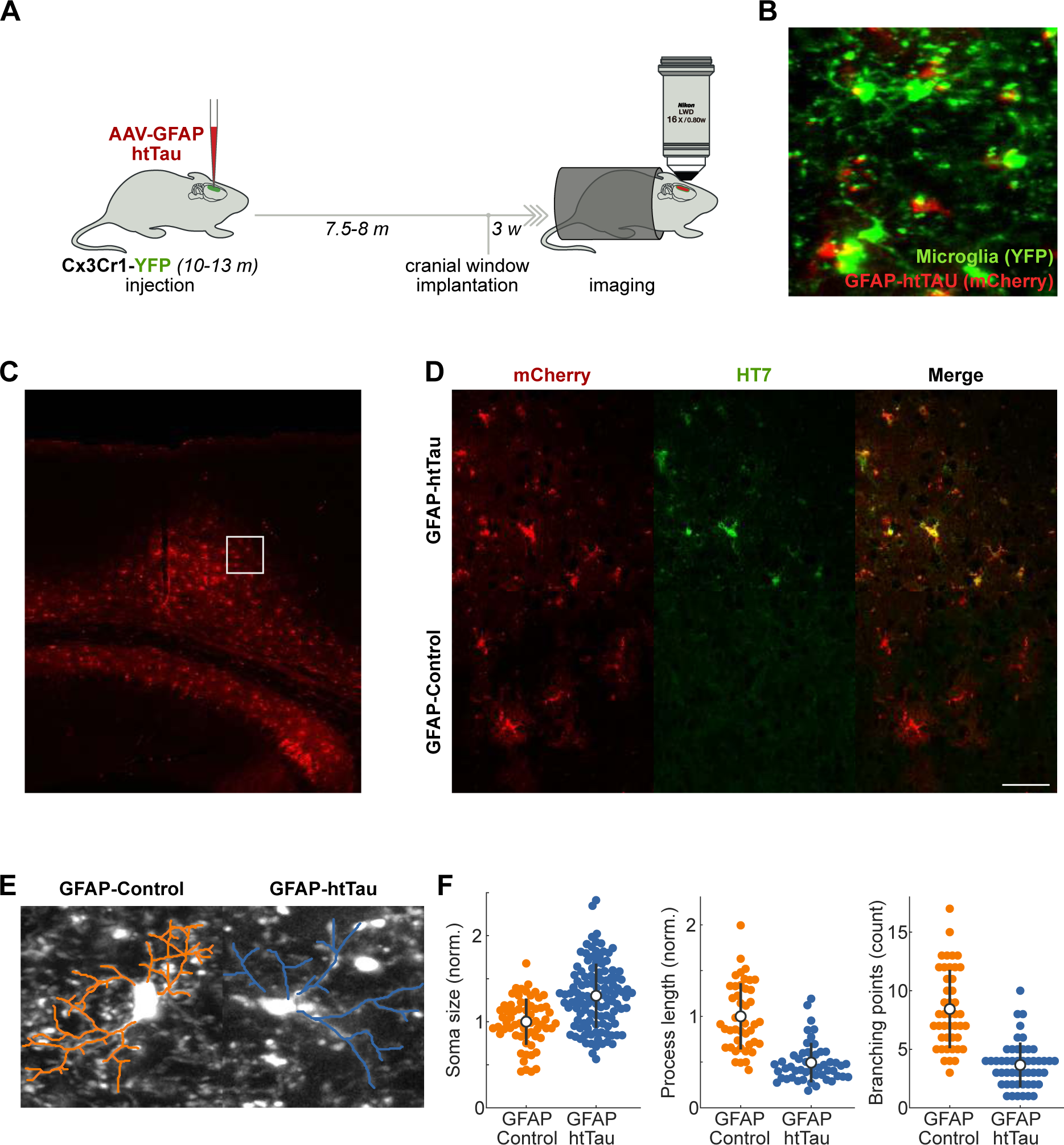
Microglial morphology changed in cortical areas with astrocytic tau pathology. **(A)** AAV-GFAP-htTau was injected into the cortex of Cx3Cr1-YFP mice and microglia were imaged 8 months after the injection. **(B)** Example maximum projection of a z-stack showing cortical co-expression of AAV-GFAP-htTau (mCherry, red) and microglia (CX3CR1-YFP, green) as visualized with *in vivo* 2-photon imaging. **(C)** Example of a brain slice of a CX3CR1-YFP mouse intra-cortically injected with AAV-GFAP-htTau. The small square corresponds to the area shown in panel D. Scale bar shows 50 μm. **(D)** Expression of human tau (HT7, green) in the cortex of CX3CR1-YFP mice injected either with AAV-GFAP-htTau or AAV-GFAP-Control (mCherry, red). Scale bar shows 50 μm. **(E)** Example of morphological analyses of microglial processes in a mouse injected with AAV-GFAP-Control (left) or AAV-GFAP-htTau (right). **(F)** Microglial soma size was larger (on average), while the total process length and the number of branches were smaller in GFAP-htTau mice compared to GFAP-Control mice. *Soma size:* GFAP-htTau 1594.46±457.45, CI=1511.23–1678.74, n=113; GFAP-Control 1225.48±329.34, CI=1141.17–1301.70, n=63; *Process length:* GFAP-htTau 257.32±111.85, CI=1511.23 – 1678.74, n=50; GFAP-Control 520.63±188.60, CI=469.18 – 579.88, n=44; *Branching:* GFAP-htTau 3.66±1.92, CI=3.20–4.26, n=50; GFAP-Control 8.43±3.34, CI=7.52–9.45, n=44. Empty circles and error bars show means and standard deviations.

We analyzed the morphology of microglia by analyzing their process length, process branching, and soma size in mice injected with AAV-GFAP-hTau or AAV-GFAP-Control **(Figure 4E)**. Estimated soma sizes and process lengths were normalized to the means of their respective control populations. Microglia in the vicinity of truncated tau-positive astrocytes had increased soma size (GFAP-htTau 1.30±0.37; GFAP-Control 1.00±0.27; p<0.001). The main microglial process with all side processes was shorter in animals injected with AAV-GFAP-htTau compared to controls (GFAP-htTau 0.49±0.21; GFAP-Control 1.00±0.36; p<0.001), and the number of branchpoints on the main process was lower as well (GFAP-htTau 3.7±1.92; GFAP-Control 8.4±3.34; p<0.001) (Figure 4F).

### Astrocytic tau pathology was not associated with alterations in spontaneous neuronal activity

Astrocytes are thought to play an important role in preventing neuronal hyperexcitability and excitotoxicity. We hypothesized that accumulation of truncated tau in astrocytes leads to impairments in synaptic function, which in turn could result in alterations of neuronal activity. To address this question, we used AAV-GFAP-htTau or AAV-GFAP-Control to express human truncated tau in the cortex of C57bl/6 mice and then expressed calcium indicator GCaMP6f in cortical neurons **(Figure 5A,B**, see Methods for details). We imaged spontaneous activity of pyramidal neurons surrounding truncated tau-expressing astrocytes (layer 2/3) in awake head-fixed mice five months after the initial injection **(Figure 5C,D)**. We did not, however, detect changes in any parameters we evaluated, such as the frequency distribution of activity transients or the overall neuronal activity, in animals injected with AAV-GFAP-htTau compared to controls (**Figure 5E,F).** Astrocytic tau inclusions in the early stages of astrocytic tau pathology, thus, did not lead to alterations in spontaneous neuronal activity.

**Figure 5.**
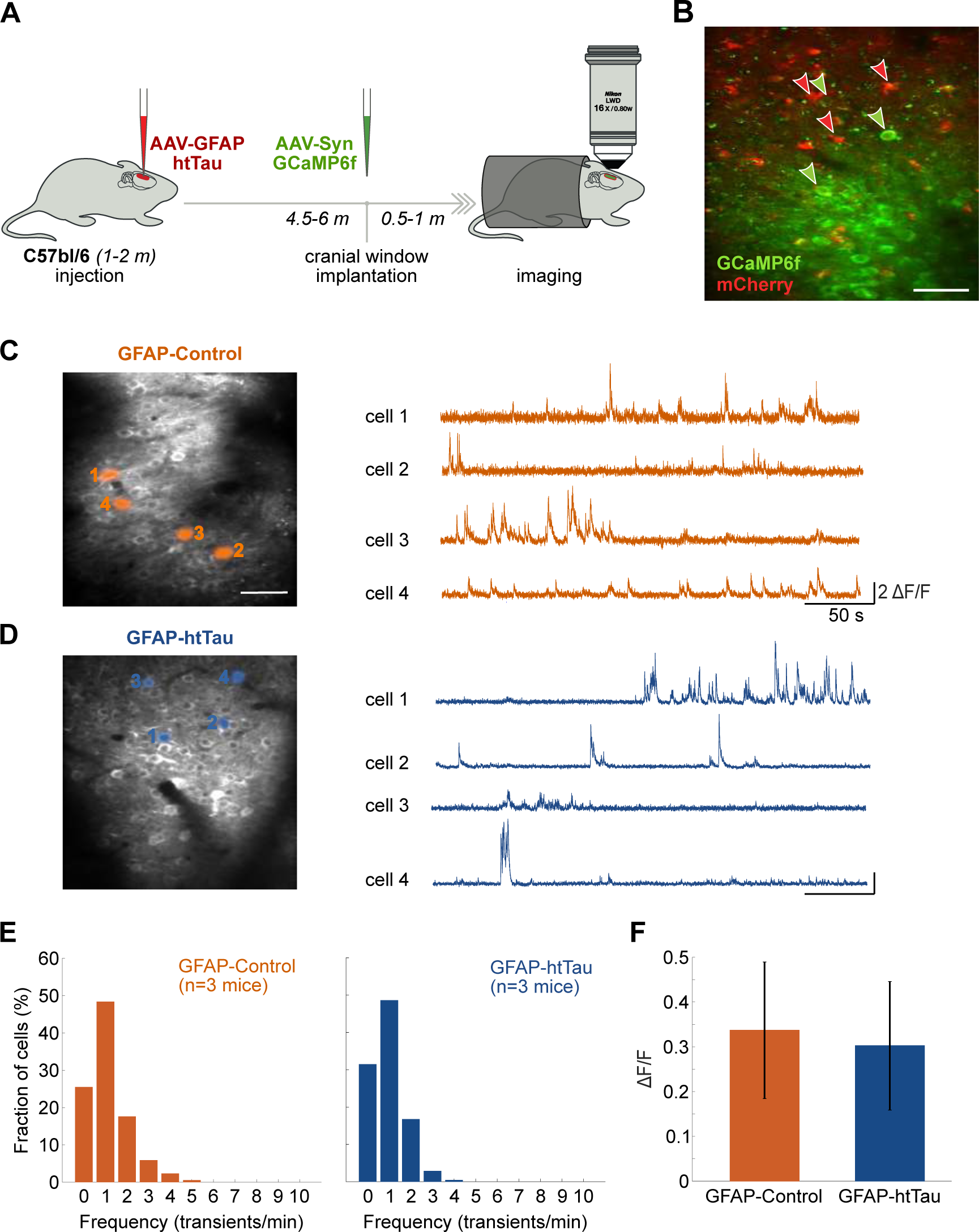
No alterations of spontaneous neuronal activity were detected in cortical areas with astrocytic tau pathology. **(A)** C57Bl/6 mice were injected with AAV-GFAP-htTau or AAV-GFAP-Control, followed by AAV-Syn-GCaMP6f, and the activity of cortical neurons was imaged using in vivo two-photon imaging. **(B)** Cortical co-expression of mCherry (GFAP-htTau) in astrocytes (red) and GCaMP6f in neurons (green) in vivo. Scale bar shows 100 μm. **(C, D)** Example two-photon average fluorescence GCaMP6f-expressing 2/3 layer neurons of AAV-GFAP-Control (C) or AAV-GFAP-htTAu (D) injected wild-type mice and corresponding neuronal activity of four example neurons normalized to baseline. Scale bar shows 100 μm. **(E)** Frequency distribution of Ca^2+^ transients of neurons recorded in GFAP-Control mice (orange, 401 neurons in 3 mice) and GFAP-htTau mice (blue, 889 neurons in 3 mice). **(F)** Average firing activity (ΔF/F) was the same for neurons recorded in GFAP-Control (orange, n = 3 mice) and GFAP-htTau (blue, n = 3 mice) mice. Error bars show standard deviations.

## Discussion

Here we have described an AAV-based model of astrocyte-specific tau pathology that can be used to induce astrocytic human truncated tau-mediated pathology in the brain areas of interest, such as the hippocampus or selected cortical regions. Transduced cells with astrocytic morphology were positive for human truncated tau, and after more than five months after injection we observed accumulation of soluble phosphorylated tau in transduced astrocytes. Astrocytic tau pathology was accompanied by alterations in microglial morphology. We did not, however, detect any cognitive impairment or alterations in spontaneous neuronal activity. The results indicate that early-stage astrocytic tau pathology is associated with microglial alterations but perhaps not yet with neuronal dysfunction. In human patient brain material, it is challenging to disentangle the contribution of different disease processes. Based on a model of astrocyte-specific tauopathy, our findings suggest that microglial abnormalities are linked to the presence of astrocytic tau inclusions in primary tauopathies.

Microglia are dynamic cells that constantly scan the brain parenchyma and are sensitive to signs of abnormalities and damage [12, 51]. Additionally, there is a strong bi-directional cross-talk between microglia and astrocytes, and alterations in the signaling pathways have been linked to neuronal tau pathology [68]. Microglia were also shown to alter their morphology and process dynamics in response to neurons with hyperphosphorylated tau in zebrafish [21]. Further, astrocytic tau pathology in PSP was associated with an immune network enriched in microglial genes [1]. Microglial activation and dystrophic microglia have been observed in animals modes of neuronal tau pathology and neuronal tauopathies such as AD [68]. Microgliosis has also been observed in conditions with astrocytic tau pathology, such as PSP, CBD, and PiD [4, 76]. Traditionally, it has been challenging to link microglial alterations to astrocytic pathology conclusively due to the presence of neuronal tau pathology and other pathological processes (e.g. neurodegeneration, co-pathologies). We employed *in vivo* imaging to study the effects of astrocytic tau pathology on microglial morphology. We observed a striking reduction in microglia process complexity and a mild increase in soma size, comparable to what has been observed using *in vivo* imaging of aged microglia in mice [22], and reminiscent of the dystrophic microglia observed in the vicinity of neuronal tau pathology in AD [68]. Previously, it has been described that astrocytic tau pathology leads to cellular degeneration rather than reactive astrogliosis [47, 66]. It is unclear, however, what causes microglial alterations in the context of tau pathology. Although microglia are able to phagocytose extracellular tau [5, 39, 41], tau inclusions are generally not found in microglia of primary tauopathies. In contrast to AD, extracellular tau is not increased in PSP, and no seed-competent tau was detected in primary microglia derived from PSP patients [25, 70]. It is, therefore, possible that astrocytic tau pathology leads to altered signaling between microglia and affected astrocytes rather than tau having a direct effect on microglia.

Astrocytic processes form intimate connections with neuronal synapses, forming functional units referred to as the ‘tripartite synapse’. A single cortical astrocyte can connect to up to 100.000 synapses in mice and 2 million synapses in humans [2]. Approximately 90% of cortical synapses are ensheathed by astrocytic processes in a dynamic process which is at least partly dependent on neuronal activity [14]. Astrocytic tau pathology in the spinal cord leads to motor problems in mice as early as four months of age, despite the fact that phospho-tau pathology was only observed in 95% of 24-month old animals. [11, 17]. These later stages were also associated with axonal pathology and elin breakdown in the absence of neuron loss [17]. In addition, lentiviral-induced expression of 3R tau in astrocytes was reported to lead to cognitive deficits [59]. These studies suggest that even very early stages of astrocytic tau pathology can lead to impairments in neuronal function and associated network dysfunction. Indeed, astrocytic tau pathology leads to loss of astrocytic glutamate transporter GLT-1 in a transgenic mouse model and patients with CBD or GGT-associated astrocytic tau pathology [11, 15]. This transporter mediates 95% of glutamate uptake in the brain [62, 65]. Astrocytic tau pathology in patients was also associated with synapse loss [8]. We therefore hypothesized that pathological accumulation of truncated tau in astrocytes might lead to alterations in neuronal activity. However, we did not detect cognitive impairments or neuronal network deficits in mice injected with AAV-GFAP-htTau at five months post-injection. It is possible that astrocytic tau pathology at this timepoint will lead to more subtle alterations in neuronal network dysfunction that does not present as overall functional impairment. It also possible that astrocytic tau pathology leads to only mild clinical symptoms. Indeed, incidental cases of CBD were asymptomatic despite having astrocytic plaques [36, 42, 46], and CBD cases with a more aggressive disease course contained more neuronal tau aggregates relative to astrocytic plaques [35].

AAV-based models represent a promising advance in studying primary tauopathies, but AAV vectors could be modified to improve model parameters. The mild off-target expression of truncated tau in neurons we observed could be eliminated and cell-type specificity improved further by using a different AAV serotype. For example, AAV5 serotype displays strong astrocytic selectivity, even in the absence of an astrocyte-specific promotor [53]. We have restricted our analysis to regions with high astrocyte specificity such as CA1 and the cortex, but behavioral testing might be challenging to interpret in the presence of apparent off-target labelling in anatomical areas like the dentate gyrus.

The astrocytic tau pathology in the presented AAV model was relatively mild, with sparser populations of astrocytes with intracellular accumulation of phosphorylated tau and low amounts of insoluble tau. Although widely used, the GFAP promotor was reported to cause relatively weak expression and might therefore not be sufficient to induce widespread accumulation of tau pathology at timepoints suitable for *in* vivo research [26]. Other astrocyte-specific promotors could be considered for future studies (e.g. ALDH1L1, GLAST) [45]. Alternatively, an enhanced GFAP promotor (using e.g. upstream insertion of CMV enhancers) might be employed to drive stronger expression [26]. Furthermore, the addition of multiple frontotemporal dementia-related *MAPT* mutations to overexpressed tau protein was reported to dramatically increase the fibrillization in AAV-models of neuronal tau pathology [10]. These mutations could be combined with non-mutated truncated tau construct and might lead to more widespread accumulation of aggregated tau in astrocytes and potentially more pronounced functional deficits.

We have developed a versatile animal model of astrocytic tau pathology using an AAV vector expressing human truncated tau protein under the control of astrocytic GFAP promotor. Such models open the door to studying the functional consequences of cell type-specific tau pathology. The origins of astrocytic tau pathology are unknown, but astrocytes have been shown to phagocytose extracellular tau and spread it to neurons or glial cells [43, 55, 71, 75]. AAV models that co-express fluorophores have previously been used to study spreading of neuronal tau pathology [57, 69, 72–74]. Interestingly, these models can be used to study spreading of human tau from neurons to astrocytes and microglia [5, 44, 57]. AAV-GFAP-htTau can be used to study the mechanisms of tau spreading from astrocytes to other cell types, which may underlie the progression of astrocytic tau pathology in conditions like ARTAG, PSP, and even AD [16, 30, 31]. We suggest that the presented AAV vector and related variants form an important toolkit for studying the functional consequences of astrocytic tau pathology in vivo.

## Acknowledgments

We would like to thank Andrej Kováč, Petra Majerová, Tomáš Smolek, Kristína Šešerová, Marian Horváth, and Peter Szalay for assistance with experiments.

The study was supported by a grant from the Slovak Research and Development Agency APVV-19-0585 to TH. TV was also supported by the European Union’s Horizon 2020 research and innovation programme under the Marie Skłodowska-Curie grant agreement No 676144 (Synaptic Dysfunction in Alzheimer Disease, SyDAD).

We would like to thank the Netherlands Brain Bank and the patients for donating the brain material.

## References

1. Allen M, Wang X, Serie DJ, Strickland SL, Burgess JD, Koga S, Younkin CS, Nguyen TT, Malphrus KG, Lincoln SJ, Alamprese M, Zhu K, Chang R, Carrasquillo MM, Kouri N, Murray ME, Reddy JS, Funk C, Price ND, Golde TE, Younkin SG, Asmann YW, Crook JE, Dickson DW, Ertekin-Taner N (2018) Divergent brain gene expression patterns associate with distinct cell-specific tau neuropathology traits in progressive supranuclear palsy. Acta Neuropathol 136:709–727. doi: 10.1007/s00401-018-1900-5

2. Allen NJ, Eroglu C (2017) Cell Biology of Astrocyte-Synapse Interactions. Neuron 96:697–708. doi: 10.1016/j.neuron.2017.09.056

3. Allen NJ, Lyons DA (2018) Glia as architects of central nervous system formation and function. Science 362:181–185. doi: 10.1126/science.aat0473

4. Alster P, Madetko N, Koziorowski D, Friedman A (2020) Microglial Activation and Inflammation as a Factor in the Pathogenesis of Progressive Supranuclear Palsy (PSP). Front Neurosci 14:893. doi: 10.3389/fnins.2020.00893

5. Asai H, Ikezu S, Tsunoda S, Medalla M, Luebke J, Haydar T, Wolozin B, Butovsky O, Kugler S, Ikezu T (2015) Depletion of microglia and inhibition of exosome synthesis halt tau propagation. Nat Neurosci 18:1584–1593. doi: 10.1038/nn.4132

6. Baskota SU, Lopez OL, Greenamyre JT, Kofler J (2019) Spectrum of tau pathologies in Huntington’s disease. Lab Invest 99:1068–1077. doi: 10.1038/s41374-018-0166-9

7. Boluda S, Iba M, Zhang B, Raible KM, Lee VM-Y, Trojanowski JQ (2015) Differential induction and spread of tau pathology in young PS19 tau transgenic mice following intracerebral injections of pathological tau from Alzheimer’s disease or corticobasal degeneration brains. Acta Neuropathol 129:221–237. doi: 10.1007/s00401-014-1373-0

8. Briel N, Pratsch K, Roeber S, Arzberger T, Herms J (2020) Contribution of the astrocytic tau pathology to synapse loss in progressive supranuclear palsy and corticobasal degeneration. Brain Pathol e12914. doi: 10.1111/bpa.12914

9. Clavaguera F, Akatsu H, Fraser G, Crowther RA, Frank S, Hench J, Probst A, Winkler DT, Reichwald J, Staufenbiel M, Ghetti B, Goedert M, Tolnay M (2013) Brain homogenates from human tauopathies induce tau inclusions in mouse brain. Proc Natl Acad Sci U S A 110:9535– 9540. doi: 10.1073/pnas.1301175110

10. Croft CL, Cruz PE, Ryu DH, Ceballos-Diaz C, Strang KH, Woody BM, Lin W-L, Deture M, Rodriguez-Lebron E, Dickson DW, Chakrabarty P, Levites Y, Giasson BI, Golde TE (2019) rAAV-based brain slice culture models of Alzheimer’s and Parkinson’s disease inclusion pathologies. J Exp Med 216:539–555. doi: 10.1084/jem.20182184

11. Dabir D V, Robinson MB, Swanson E, Zhang B, Trojanowski JQ, Lee VM-Y, Forman MS (2006) Impaired glutamate transport in a mouse model of tau pathology in astrocytes. J Neurosci 26:644–654. doi: 10.1523/JNEUROSCI.3861-05.2006

12. Davalos D, Grutzendler J, Yang G, Kim J V, Zuo Y, Jung S, Littman DR, Dustin ML, Gan W-B (2005) ATP mediates rapid microglial response to local brain injury in vivo. Nat Neurosci 8:752–758. doi: 10.1038/nn1472

13. Dawson HN, Cantillana V, Chen L, Vitek MP (2007) The tau N279K exon 10 splicing mutation recapitulates frontotemporal dementia and parkinsonism linked to chromosome 17 tauopathy in a mouse model. J Neurosci 27:9155–9168. doi: 10.1523/JNEUROSCI.5492-06.2007

14. Farhy-Tselnicker I, Allen NJ (2018) Astrocytes, neurons, synapses: a tripartite view on cortical circuit development. Neural Dev 13:7. doi: 10.1186/s13064-018-0104-y

15. Ferrer I, Andres-Benito P, Zelaya MV, Aguirre MEE, Carmona M, Ausin K, Lachen-Montes M, Fernandez-Irigoyen J, Santamaria E, Del Rio JA (2020) Familial globular glial tauopathy linked to MAPT mutations: molecular neuropathology and seeding capacity of a prototypical mixed neuronal and glial tauopathy. Acta Neuropathol. doi: 10.1007/s00401-019-02122-9

16. Fleeman RM, Proctor EA (2021) Astrocytic Propagation of Tau in the Context of Alzheimer’s Disease. Front Cell Neurosci 15:645233. doi: 10.3389/fncel.2021.645233

17. Forman MS, Lal D, Zhang B, Dabir D V, Swanson E, Lee VM-Y, Trojanowski JQ (2005) Transgenic Mouse Model of Tau Pathology in Astrocytes Leading to Nervous System Degeneration. J Neurosci 25:3539 LP – 3550

18. Goedert M, Spillantini MG, Potier MC, Ulrich J, Crowther RA (1989) Cloning and sequencing of the cDNA encoding an isoform of microtubule-associated protein tau containing four tandem repeats: differential expression of tau protein mRNAs in human brain. EMBO J 8:393–399

19. Hallmann A-L, Arauzo-Bravo MJ, Mavrommatis L, Ehrlich M, Ropke A, Brockhaus J, Missler M, Sterneckert J, Scholer HR, Kuhlmann T, Zaehres H, Hargus G (2017) Astrocyte pathology in a human neural stem cell model of frontotemporal dementia caused by mutant TAU protein. Sci Rep 7:42991. doi: 10.1038/srep42991

20. Hammond SL, Leek AN, Richman EH, Tjalkens RB (2017) Cellular selectivity of AAV serotypes for gene delivery in neurons and astrocytes by neonatal intracerebroventricular injection. PLoS One 12:e0188830–e0188830. doi: 10.1371/journal.pone.0188830

21. Hassan-Abdi R, Brenet A, Bennis M, Yanicostas C, Soussi-Yanicostas N (2019) Neurons expressing pathological Tau protein trigger dramatic changes in microglial morphology and dynamics. bioRxiv 731901. doi: 10.1101/731901

22. Hefendehl JK, Neher JJ, Suhs RB, Kohsaka S, Skodras A, Jucker M (2014) Homeostatic and injury-induced microglia behavior in the aging brain. Aging Cell 13:60–69. doi: 10.1111/acel.12149

23. Higuchi M, Ishihara T, Zhang B, Hong M, Andreadis A, Trojanowski J, Lee VM-Y (2002) Transgenic mouse model of tauopathies with glial pathology and nervous system degeneration. Neuron 35:433–446

24. Hoglinger GU, Respondek G, Kovacs GG (2018) New classification of tauopathies. Rev Neurol (Paris). doi: 10.1016/j.neurol.2018.07.001

25. Hopp SC, Lin Y, Oakley D, Roe AD, DeVos SL, Hanlon D, Hyman BT (2018) The role of microglia in processing and spreading of bioactive tau seeds in Alzheimer’s disease. J Neuroinflammation 15:269. doi: 10.1186/s12974-018-1309-z

26. Joshi CR, Raghavan V, Vijayaraghavalu S, Gao Y, Saraswathy M, Labhasetwar V, Ghorpade A (2018) Reaching for the Stars in the Brain: Polymer-Mediated Gene Delivery to Human Astrocytes. Mol Ther Nucleic Acids 12:645–657. doi: 10.1016/j.omtn.2018.06.009

27. Kahlson MA, Colodner KJ (2015) Glial Tau Pathology in Tauopathies: Functional Consequences. J Exp Neurosci 9:43–50. doi: 10.4137/JEN.S25515

28. Kovacs GG (2020) Astroglia and Tau: New Perspectives. Front. Aging Neurosci. 12:96

29. Kovacs GG, Lee VM, Trojanowski JQ (2017) Protein astrogliopathies in human neurodegenerative diseases and aging. Brain Pathol 27:675–690. doi: 10.1111/bpa.12536

30. Kovacs GG, Lukic MJ, Irwin DJ, Arzberger T, Respondek G, Lee EB, Coughlin D, Giese A, Grossman M, Kurz C, McMillan CT, Gelpi E, Compta Y, van Swieten JC, Laat LD, Troakes C, Al-Sarraj S, Robinson JL, Roeber S, Xie SX, Lee VM-Y, Trojanowski JQ, Höglinger GU (2020) Distribution patterns of tau pathology in progressive supranuclear palsy. Acta Neuropathol. doi: 10.1007/s00401-020-02158-2

31. Kovacs GG, Xie SX, Robinson JL, Lee EB, Smith DH, Schuck T, Lee VM-Y, Trojanowski JQ (2018) Sequential stages and distribution patterns of aging-related tau astrogliopathy (ARTAG) in the human brain. Acta Neuropathol Commun 6:50. doi: 10.1186/s40478-018-0552-y

32. Kyrargyri V, Attwell D, Jolivet RB, Madry C (2019) Analysis of Signaling Mechanisms Regulating Microglial Process Movement. Methods Mol Biol 2034:191–205. doi: 10.1007/978-1-4939-9658-2_14

33. Letellier M, Park YK, Chater TE, Chipman PH, Gautam SG, Oshima-Takago T, Goda Y (2016) Astrocytes regulate heterogeneity of presynaptic strengths in hippocampal networks. Proc Natl Acad Sci U S A 113:E2685–E2694. doi: 10.1073/pnas.1523717113

34. Lin W-L, Lewis J, Yen S-H, Hutton M, Dickson DW (2003) Filamentous tau in oligodendrocytes and astrocytes of transgenic mice expressing the human tau isoform with the P301L mutation. Am J Pathol 162:213–218. doi: 10.1016/S0002-9440(10)63812-6

35. Ling H, Gelpi E, Davey K, Jaunmuktane Z, Mok KY, Jabbari E, Simone R, R’Bibo L, Brandner S, Ellis MJ, Attems J, Mann D, Halliday GM, Al-Sarraj S, Hedreen J, Ironside JW, Kovacs GG, Kovari E, Love S, Vonsattel JPG, Allinson KSJ, Hansen D, Bradshaw T, Seto-Salvia N, Wray S, de Silva R, Morris HR, Warner TT, Hardy J, Holton JL, Revesz T (2020) Fulminant corticobasal degeneration: a distinct variant with predominant neuronal tau aggregates. Acta Neuropathol. doi: 10.1007/s00401-019-02119-4

36. Ling H, Kovacs GG, Vonsattel JPG, Davey K, Mok KY, Hardy J, Morris HR, Warner TT, Holton JL, Revesz T (2016) Astrogliopathy predominates the earliest stage of corticobasal degeneration pathology. Brain 139:3237–3252. doi: 10.1093/brain/aww256

37. Longair MH, Baker DA, Armstrong JD (2011) Simple Neurite Tracer: open source software for reconstruction, visualization and analysis of neuronal processes. Bioinformatics 27:2453– 2454.doi: 10.1093/bioinformatics/btr390

38. Lopes G, Bonacchi N, Frazão J, Neto JP, Atallah B V, Soares S, Moreira L, Matias S, Itskov PM, Correia PA, Medina RE, Calcaterra L, Dreosti E, Paton JJ, Kampff AR (2015) Bonsai: an event-based framework for processing and controlling data streams. Front. Neuroinformatics 9:7

39. Luo W, Liu W, Hu X, Hanna M, Caravaca A, Paul SM (2015) Microglial internalization and degradation of pathological tau is enhanced by an anti-tau monoclonal antibody. Sci Rep 5:11161. doi: 10.1038/srep11161

40. Madry C, Kyrargyri V, Arancibia-Carcamo IL, Jolivet R, Kohsaka S, Bryan RM, Attwell D (2018) Microglial Ramification, Surveillance, and Interleukin-1beta Release Are Regulated by the Two-Pore Domain K(+) Channel THIK-1. Neuron 97:299-312.e6. doi: 10.1016/j.neuron.2017.12.002

41. Majerova P, Zilkova M, Kazmerova Z, Kovac A, Paholikova K, Kovacech B, Zilka N, Novak M (2014) Microglia display modest phagocytic capacity for extracellular tau oligomers. J Neuroinflammation 11:161. doi: 10.1186/s12974-014-0161-z

42. Martinez-Maldonado A, Luna-Munoz J, Ferrer I (2016) Incidental corticobasal degeneration. Neuropathol. Appl. Neurobiol. 42:659–663

43. Martini-Stoica H, Cole AL, Swartzlander DB, Chen F, Wan Y-W, Bajaj L, Bader DA, Lee VMY, Trojanowski JQ, Liu Z, Sardiello M, Zheng H (2018) TFEB enhances astroglial uptake of extracellular tau species and reduces tau spreading. J Exp Med 215:2355–2377. doi: 10.1084/jem.20172158

44. Maté de Gérando A, d’Orange M, Augustin E, Joséphine C, Aurégan G, Gaudin-Guérif M, Guillermier M, Hérard A-S, Stimmer L, Petit F, Gipchtein P, Jan C, Escartin C, Selingue E, Carvalho K, Blum D, Brouillet E, Hantraye P, Gaillard M-C, Bonvento G, Bemelmans A-P, Cambon K (2021) Neuronal tau species transfer to astrocytes and induce their loss according to tau aggregation state. Brain. doi: 10.1093/brain/awab011

45. Merienne N, Le Douce J, Faivre E, Déglon N, Bonvento G (2013) Efficient gene delivery and selective transduction of astrocytes in the mammalian brain using viral vectors. Front. Cell. Neurosci. 7:106

46. Milenkovic I, Kovacs GG (2013) Incidental corticobasal degeneration in a 76-year-old woman. Clin. Neuropathol. 32:69–72

47. Motoi Y, Takanashi M, Itaya M, Ikeda K, Mizuno Y, Mori H (2004) Glial localization of four-repeat tau in atypical progressive supranuclear palsy. Neuropathology 24:60–65. doi: 10.1111/j.1440-1789.2003.00529.x

48. Munoz DG, Woulfe J, Kertesz A (2007) Argyrophilic thorny astrocyte clusters in association with Alzheimer’s disease pathology in possible primary progressive aphasia. Acta Neuropathol 114:347–357. doi: 10.1007/s00401-007-0266-x

49. Narasimhan S, Changolkar L, Riddle DM, Kats A, Stieber A, Weitzman SA, Zhang B, Li Z, Roberson ED, Trojanowski JQ, Lee VMY (2020) Human tau pathology transmits glial tau aggregates in the absence of neuronal tau. J Exp Med 217. doi: 10.1084/jem.20190783

50. Nimmerjahn A (2012) Optical window preparation for two-photon imaging of microglia in mice. Cold Spring Harb Protoc 2012. doi: 10.1101/pdb.prot069286

51. Nimmerjahn A, Kirchhoff F, Helmchen F (2005) Resting Microglial Cells Are Highly Dynamic Surveillants of Brain Parenchyma in Vivo. Science (80-) 308:1314 LP – 1318

52. Nolan A, De Paula Franca Resende E, Petersen C, Neylan K, Spina S, Huang E, Seeley W, Miller Z, Grinberg LT (2019) Astrocytic Tau Deposition Is Frequent in Typical and Atypical Alzheimer Disease Presentations. J Neuropathol Exp Neurol. doi: 10.1093/jnen/nlz094

53. Ortinski PI, Dong J, Mungenast A, Yue C, Takano H, Watson DJ, Haydon PG, Coulter DA (2010) Selective induction of astrocytic gliosis generates deficits in neuronal inhibition. Nat Neurosci 13:584–591. doi: 10.1038/nn.2535

54. Pachitariu M, Stringer C, Dipoppa M, Schröder S, Rossi LF, Dalgleish H, Carandini M, Harris KD (2017) Suite2p: beyond 10,000 neurons with standard two-photon microscopy. bioRxiv 61507. doi: 10.1101/061507

55. Perea JR, Lopez E, Diez-Ballesteros JC, Avila J, Hernandez F, Bolos M (2019) Extracellular Monomeric Tau Is Internalized by Astrocytes. Front Neurosci 13:442. doi: 10.3389/fnins.2019.00442

56. Pologruto TA, Sabatini BL, Svoboda K (2003) ScanImage: flexible software for operating laser scanning microscopes. Biomed Eng Online 2:13. doi: 10.1186/1475-925X-2-13

57. Rauch JN, Luna G, Guzman E, Audouard M, Challis C, Sibih YE, Leshuk C, Hernandez I, Wegmann S, Hyman BT, Gradinaru V, Kampmann M, Kosik KS (2020) LRP1 is a master regulator of tau uptake and spread. Nature. doi: 10.1038/s41586-020-2156-5

58. Resende E de PF, Nolan AL, Petersen C, Ehrenberg AJ, Spina S, Allen IE, Rosen HJ, Kramer J, Miller BL, Seeley WW, Gorno-Tempini ML, Miller Z, Grinberg LT (2020) Language and spatial dysfunction in Alzheimer disease with white matter thorn-shaped astrocytes: Astrocytic tau, cognitive function, and Alzheimer disease. Neurology. doi: 10.1212/WNL.0000000000008937

59. Richetin K, Steullet P, Pachoud M, Perbet R, Parietti E, Maheswaran M, Eddarkaoui S, Bégard S, Pythoud C, Rey M, Caillierez R, Q Do K, Halliez S, Bezzi P, Buée L, Leuba G, Colin M, Toni N, Déglon N (2020) Tau accumulation in astrocytes of the dentate gyrus induces neuronal dysfunction and memory deficits in Alzheimer’s disease. Nat Neurosci. doi: 10.1038/s41593-020-00728-x

60. Robinson JL, Corrada MM, Kovacs GG, Dominique M, Caswell C, Xie SX, Lee VM-Y, Kawas CH, Trojanowski JQ (2018) Non-Alzheimer’s contributions to dementia and cognitive resilience in The 90+ Study. Acta Neuropathol 136:377–388. doi: 10.1007/s00401-018-1872-5

61. Robinson JL, Yan N, Caswell C, Xie SX, Suh E, Van Deerlin VM, Gibbons G, Irwin DJ, Grossman M, Lee EB, Lee VM-Y, Miller B, Trojanowski JQ (2020) Primary Tau Pathology, Not Copathology, Correlates With Clinical Symptoms in PSP and CBD. J Neuropathol Exp Neurol 79:296–304. doi: 10.1093/jnen/nlz141

62. Rothstein JD, Dykes-Hoberg M, Pardo CA, Bristol LA, Jin L, Kuncl RW, Kanai Y, Hediger MA, Wang Y, Schielke JP, Welty DF (1996) Knockout of glutamate transporters reveals a major role for astroglial transport in excitotoxicity and clearance of glutamate. Neuron 16:675–686

63. Santacruz K, Lewis J, Spires T, Paulson J, Kotilinek L, Ingelsson M, Guimaraes A, DeTure M, Ramsden M, McGowan E, Forster C, Yue M, Orne J, Janus C, Mariash A, Kuskowski M, Hyman B, Hutton M, Ashe KH (2005) Tau suppression in a neurodegenerative mouse model improves memory function. Science 309:476–481. doi: 10.1126/science.1113694

64. Sidoryk-Wegrzynowicz M, Struzynska L (2019) Astroglial contribution to tau-dependent neurodegeneration. Biochem J 476:3493–3504. doi: 10.1042/BCJ20190506

65. Tanaka K, Watase K, Manabe T, Yamada K, Watanabe M, Takahashi K, Iwama H, Nishikawa T, Ichihara N, Kikuchi T, Okuyama S, Kawashima N, Hori S, Takimoto M, Wada K (1997) Epilepsy and Exacerbation of Brain Injury in Mice Lacking the Glutamate Transporter GLT-1. Science (80-) 276:1699 LP – 1702. doi: 10.1126/science.276.5319.1699

66. Togo T, Dickson DW (2002) Tau accumulation in astrocytes in progressive supranuclear palsy is a degenerative rather than a reactive process. Acta Neuropathol 104:398–402. doi: 10.1007/s00401-002-0569-x

67. Vogels T, Leuzy A, Cicognola C, Ashton NJ, Smolek T, Novak M, Blennow K, Zetterberg H, Hromadka T, Zilka N, Schöll M (2020) Propagation of Tau Pathology: Integrating Insights From Postmortem and In Vivo Studies. Biol Psychiatry 87:808–818. doi: 10.1016/j.biopsych.2019.09.019

68. Vogels T, Murgoci A-N, Hromadka T (2019) Intersection of pathological tau and microglia at the synapse. Acta Neuropathol Commun 7:109. doi: 10.1186/s40478-019-0754-y

69. Vogels T, Vargová G, Brezováková V, Quint WH, Hromádka T (2020) Viral Delivery of Non-Mutated Human Truncated Tau to Neurons Recapitulates Key Features of Human Tauopathy in Wild-Type Mice. J Alzheimers Dis. doi: 10.3233/JAD-200047

70. Wagshal D, Sankaranarayanan S, Guss V, Hall T, Berisha F, Lobach I, Karydas A, Voltarelli L, Scherling C, Heuer H, Tartaglia MC, Miller Z, Coppola G, Ahlijanian M, Soares H, Kramer JH, Rabinovici GD, Rosen HJ, Miller BL, Meredith J, Boxer AL (2015) Divergent CSF tau alterations in two common tauopathies: Alzheimer’s disease and progressive supranuclear palsy. J Neurol Neurosurg Psychiatry 86:244–250. doi: 10.1136/jnnp-2014-308004

71. Wang P, Ye Y (2020) An integrin receptor complex mediates filamentous Tau-induced activation of primary astrocytes. bioRxiv 2020.05.07.083295. doi: 10.1101/2020.05.07.083295

72. Wegmann S, Bennett RE, Amaral AS, Hyman BT (2017) Studying tau protein propagation and pathology in the mouse brain using adeno-associated viruses. Methods Cell Biol 141:307–322. doi: 10.1016/bs.mcb.2017.06.014

73. Wegmann S, Bennett RE, Delorme L, Robbins AB, Hu M, McKenzie D, Kirk MJ, Schiantarelli J, Tunio N, Amaral AC, Fan Z, Nicholls S, Hudry E, Hyman BT (2019) Experimental evidence for the age dependence of tau protein spread in the brain. Sci Adv 5:eaaw6404. doi: 10.1126/sciadv.aaw6404

74. Wegmann S, Maury EA, Kirk MJ, Saqran L, Roe A, DeVos SL, Nicholls S, Fan Z, Takeda S, Cagsal-Getkin O, William CM, Spires-Jones TL, Pitstick R, Carlson GA, Pooler AM, Hyman BT (2015) Removing endogenous tau does not prevent tau propagation yet reduces its neurotoxicity. EMBO J 34:3028–3041. doi: 10.15252/embj.201592748

75. Wiersma VI, van Ziel AM, Vazquez-Sanchez S, Nolle A, Berenjeno-Correa E, Bonaterra-Pastra A, Clavaguera F, Tolnay M, Musters RJP, van Weering JRT, Verhage M, Hoozemans JJM, Scheper W (2019) Granulovacuolar degeneration bodies are neuron-selective lysosomal structures induced by intracellular tau pathology. Acta Neuropathol. doi: 10.1007/s00401-019-02046-4

76. Woollacott IOC, Toomey CE, Strand C, Courtney R, Benson BC, Rohrer JD, Lashley T (2020) Microglial burden, activation and dystrophy patterns in frontotemporal lobar degeneration. J Neuroinflammation 17:234. doi: 10.1186/s12974-020-01907-0

77. Xu H-T, Pan F, Yang G, Gan W-B (2007) Choice of cranial window type for in vivo imaging affects dendritic spine turnover in the cortex. Nat Neurosci 10:549–551. doi: 10.1038/nn1883

78. Yeo S, Bandyopadhyay S, Messing A, Brenner M (2013) Transgenic analysis of GFAP promoter elements. Glia 61:1488–1499. doi: 10.1002/glia.22536

79. Zhang Y, Chen K, Sloan SA, Bennett ML, Scholze AR, O&#039;Keeffe S, Phatnani HP, Guarnieri P, Caneda C, Ruderisch N, Deng S, Liddelow SA, Zhang C, Daneman R, Maniatis T, Barres BA, Wu JQ (2014) An RNA-Sequencing Transcriptome and Splicing Database of Glia, Neurons, and Vascular Cells of the Cerebral Cortex. J Neurosci 34:11929 LP – 11947. doi: 10.1523/JNEUROSCI.1860-14.2014

80. Zilka N, Filipcik P, Koson P, Fialova L, Skrabana R, Zilkova M, Rolkova G, Kontsekova E, Novak M (2006) Truncated tau from sporadic Alzheimer’s disease suffices to drive neurofibrillary degeneration in vivo. FEBS Lett 580:3582–3588. doi: 10.1016/j.febslet.2006.05.029

81. Zimova I, Brezovakova V, Hromadka T, Weisova P, Cubinkova V, Valachova B, Filipcik P, Jadhav S, Smolek T, Zilka N (2016) Human Truncated Tau Induces Mature Neurofibrillary Pathology in a Mouse Model of Human Tauopathy. J Alzheimers Dis. doi: 10.3233/JAD-160347

